# AutoGVP: a dockerized workflow integrating ClinVar and InterVar germline sequence variant classification

**DOI:** 10.1101/2023.11.29.569103

**Authors:** Jung Kim, Ammar S. Naqvi, Ryan J. Corbett, Rebecca S. Kaufman, Zalman Vaksman, Miguel A. Brown, Daniel P. Miller, Saksham Phul, Zhuangzhuang Geng, Phillip B. Storm, Adam C. Resnick, Douglas R. Stewart, Jo Lynne Rokita, Sharon J. Diskin

## Abstract

**Summary:** With the increasing rates of exome and whole genome sequencing, the ability to classify large sets of germline sequencing variants using up-to-date American College of Medical Genetics – Association for Molecular Pathology (ACMG-AMP) criteria is crucial. Here, we present Automated Germline Variant Pathogenicity (AutoGVP), a tool that integrates germline variant pathogenicity annotations from ClinVar and sequence variant classifications from a modified version of InterVar (PVS1 strength adjustments, removal of PP5/BP6). This tool facilitates large-scale, clinically-focused classification of germline sequence variants in a research setting.

**Availability and Implementation:** AutoGVP is an open-source dockerized workflow implemented in R and freely available on GitHub at https://github.com/diskin-lab-chop/AutoGVP.

## 1. Introduction

As exome and whole genome sequencing become more prevalent in clinical genetic testing and population-scale biobanks become more widely accessible, tools that facilitate classification of large sets of germline variants are crucial. Tools such as InterVar (Li and Wang, 2017), CharGer (Scott *et al*., 2019), Franklin (https://franklin.genoox.com), and Varsome (Kopanos *et al*., 2019) implement the ACMG-AMP guidelines for the interpretation of germline sequence variants (Richards *et al*., 2015). However, misclassification of variant pathogenicity may occur depending on the interpretation of specific ACMG-AMP criteria set by the tool at the time of the development. For example, variants can be mis-classified by applying “full strength” of the PVS1 rule (“a null variant in a gene where loss-of-function is a known mechanism of disease”), and/or use of the PP5 rule (“reputable source recently reports variant as pathogenic, but the evidence is not available to the laboratory to perform an independent evaluation”). In 2018, ACMG-AMP recommendations for PVS1 were modified so that null variants should not be considered at the same strength, but in different categories (PVS1, PVS1_Strong, PVS1_Moderate, PVS1_Supporting) (Abou Tayoun *et al*., 2018). To address the modified PVS1 rule, Xiang et al. developed AutoPVS1 to automate PVS1 adjustments (Abou Tayoun *et al*., 2018; Xiang *et al*., 2020). Furthermore, some tools are limited in the number of variants that can be classified at once or require the purchase of a license.

InterVar reports each ACMG-AMP criterion with a score and each criterion is easily modifiable using open-source code accessible through GitHub. Concomitant with the development of these tools has been the dramatic increase in both the quantity and quality of germline variant classifications within the ClinVar database. This is due to the collaborative effort of ClinGen variant curation expert panels (Rehm *et al*., 2015) which meticulously refine, on a gene-specific basis, ACMG-AMP criteria using clinical (phenotype) information, the underlying biology of a gene (and structure of associated protein(s)), bioinformatics predictions and population genetics. It is important to note that (as of November 2023) InterVar does *not* automatically incorporate classification rules developed by ClinGen Variant Curation Expert Panels (VCEP; https://clinicalgenome.org/affiliation/vcep/). In addition, ClinVar will only have variants that have been curated and deposited. There is a need for a comprehensive pipeline that not only annotates patient germline variation deposited to ClinVar, but also those not documented in ClinVar. We created the Automated Germline Variant Pathogenicity (AutoGVP) pipeline to classify germline variant pathogenicity in a research setting through integrating both ClinVar and InterVar annotations. Uniquely, AutoGVP automatically modifies InterVar to classify variants using most current ACMG-AMP guidelines, by adjusting PVS1 strength (Rehm *et al*., 2015; Abou Tayoun *et al*., 2018), using autoPVS1 and removing the PP5 and BP6 rules (Biesecker *et al*., 2018).

Adjusting PVS1 strength is crucial since InterVar can over-interpret loss-of-function (LOF) variants (Singer-Berk *et al*., 2023) as it classifies all LOF variants that are in LOF-intolerant genes curated by InterVar as PVS1 (Li and Wang, 2017). AutoGVP can classify any number of variants in standard VCF format. The code is freely available as a dockerized workflow in GitHub (https://github.com/diskin-lab-chop/AutoGVP) (Naqvi *et al*., 2023) with options of 1) supply a VEP-annotated VCF with ANNOVAR (Wang *et al*., 2010), AutoPVS1 (Xiang *et al*., 2020), and InterVar (Li and Wang, 2017) outputs with the flexibility to update any of these and/or the provided ClinVar versions or 2) use output from a cloud-based CAVATICA preprocessing workflow (https://github.com/d3b-center/D3b-Pathogenicity-Preprocessing) and public CAVATICA app (https://cavatica.sbgenomics.com/public/apps/cavatica/apps-publisher/d3b-diskin-pathogenicity-preprocess-wf) on germline calls from the Kids First pipeline (https://github.com/kids-first/kf-germline-workflow/blob/v0.4.4/docs/GERMLINE_SNV_ANNOT_README.md, >= v0.4.4) (**Figure 1A**). Depending on sample input size, AutoGVP can be run locally, on Amazon Web Services Elastic Compute Cloud (AWS EC2), or on a high-performance computing (HPC) cluster.

**Figure 1.**
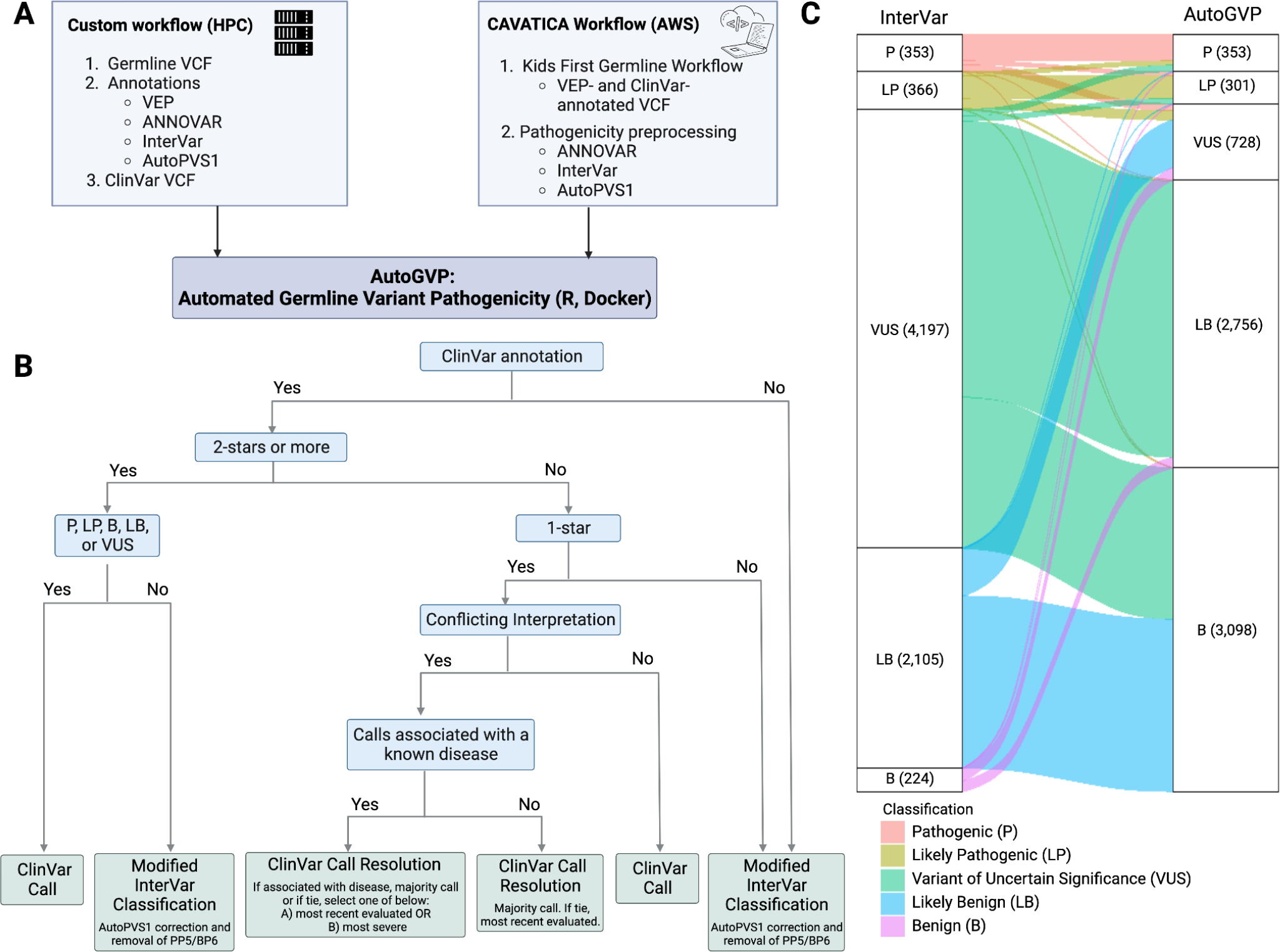
Flow diagram of AutoGVP. A) Required input files for custom or CAVATICA workflow. B) Variant classification method decision tree. C) Alluvial plot showing how InterVar classification changes in AutoGVP. The large numbers of concordant VUS (357,656), LB (20,374), and B (28,310) variants were removed for easier visualization. Numbers in parentheses represent a number of variants in that classification. Panels A and B created with BioRender.com

## 2. Material and Methods

### 2.1 Data input and preparation

#### AutoGVP is implemented in R and bash and is run within the Docker image

https://pgc-images.sbgenomics.com/naqvia/autogvp:latest. Detailed instructions on how to run AutoGVP are documented within the README of the GitHub repository. The script requires the following input files that must be supplied by the user: a VCF file with VEP annotations (*VEP.vcf), a multianno file generated by ANNOVAR run with ‘--vcfinput’ flag (*hg38_multianno.txt), an InterVar output file (*multianno.txt.intervar), and an AutoPVS1 output file (*autopvs1.txt). Additionally, it requires a ClinVar VCF file (clinvar.vcf.gz), variant summary file (variant_summary.txt.gz), and submission summary file (submission_summary.txt.gz) which can be downloaded directly from the ClinVar FTP site (https://ftp.ncbi.nlm.nih.gov/pub/clinvar/) or are stably provided through the ‘download_clinvar.sh’ script (ClinVar 7/17/2023 database). Alternative archived ClinVar version database files can also be supplied by the user. The ‘select-clinVar-submissions.R’ script generates a subset of the ClinVar database prior to running AutoGVP. Several arguments allow for customization of 1-star conflicting variant resolution. The ‘--conceptID_list’ flag takes a user-defined MedGen Concept ID list and filters variant submissions for only those associated with provided Concept IDs. As a use case, we supplied Concept IDs associated with cancer, though IDs for any disease can be used to identify P/LP variants specifically linked to the disease of interest. When conflicts cannot be resolved through consensus, users can specify whether to resolve with a Concept ID list by taking the call of the most recent submission (’--latest’) or the most severe call (’--most_severe’). Prior to running variant annotation, AutoGVP will minimally filter for PASS variants as well as any user-provided filters added in the ‘--filter_criteria’ argument (e.g., minimum variant depth, variant allele frequency). Example test files for each workflow are provided within the GitHub repository.

### 2.2 Variant annotation and classification

AutoGVP uses a hierarchical system to integrate ClinVar and InterVar variant annotations (**Figure 1B**). ClinVar variant classifications have different levels of review status represented by stars ranging from zero to four (https://www.ncbi.nlm.nih.gov/clinvar/docs/review_status/#revstat) (Rehm *et al*., 2015). AutoGVP retains the ClinVar classification for variants with two or more stars or 1-star with criteria provided by a single submitter. Variants that have 1-star with criteria provided, but have conflicting interpretations, are resolved through a multi-step decision tree with user-defined options. First, all variant submissions with “no assertion criteria provided” are excluded from the resolution process. In the absence of a Concept ID list, AutoGVP will initially try to resolve conflicts by determining a majority call, and will otherwise take the call of the most recent submission. When a Concept ID list is supplied, submissions will be filtered for only those associated with provided Concept IDs, and AutoGVP will again search for a majority call. Otherwise, conflicts will be resolved by taking the call of the most recent submission by default or if the ‘--latest’ argument is specified, will take the most severe call if ‘--most_severe’ is specified, in which order of Pathogenic (P), Likely Pathogenic (LP), Variant of Uncertain Significance (VUS), Likely Benign (LB), Benign (B) (**Figure 1B**). Variants with zero stars and variants that have non-standard P/LP/VUS/LB/B calls in ClinVar (ie. “Affects|risk_factor”, “association_not_found”, etc.) are treated as no ClinVar classification.

Variants unclassified by the ClinVar database undergo classification by InterVar, which was modified in the following two ways. First, since ClinGen recommends using the LOF PVS1 rule with varying modifications of strength (Abou Tayoun *et al*., 2018), we adjusted InterVar output using AutoPVS1 calls (Xiang *et al*., 2020). Second, since exclusion of the PP5/BP6 criterion is now recommended to prevent potential misuse and double counting (Abou Tayoun *et al*., 2018; Biesecker *et al*., 2018), we removed those two rules from InterVar. Furthermore, automated tools often use ClinVar to interpret the PP5 rule, and as a result, may capture a P/LP classification from Online Mendelian Inheritance in Man (OMIM: https://www.omim.org/) or GeneReviews (Adam *et al*., 2023) (zero stars) when in essence there is no indication of the specific variants in those two sources.

In summary, AutoGVP will generate a final call to be one of P/LP/VUS/LB/B based on the integration of the ClinVar and modified InterVar calls described above. AutoGVP modifies the InterVar classification using corrected PVS1 strength and completely removes PP5/BP6, per the most recent guidance from ClinGen. Lastly, AutoGVP will annotate variants to a single gene transcript by selecting the transcript annotation provided in AutoPVS1 file. Additional gene and transcript annotations, when present, are retained in the output column “csq_vep”. The software saves two output files per run: an abridged file with minimal columns including coordinate, gene annotation, and pathogenicity assessment information, and a full output file containing additional annotation columns from the input VCF, ANNOVAR, InterVar, and AutoPVS1 files. The current pipeline was tested using data from dbGaP phs000720.v4.p1, VEP 104, InterVar v.2.2.1, AutoPVS1 v.2.0, ClinVar database 10/28/23, and AutoGVP v0.4.2.

## 3. Results

To evaluate AutoGVP performance, we annotated 413,585 rare germline variants (gnomAD v3.1.1 non-cancer < 0.001) with 582.7MB size file from 121 individuals with rhabdomyosarcoma (dbGaP phs000720.v4.p1)(Kim *et al*., 2021) using HPC and CAVATICA. Computational time and usage to annotate above file using Concept ID with ‘--latest’ flag are for HPC, a total of 10 minutes with 0.33 CPU hours, and 36.2GB RAM were needed and for CAVATICA, a total of 9.1 minutes with 0.01 CPU hours, and 37.7GB RAM were needed. AutoGVP annotations were compared to InterVar alone (**Supplementary Table S1**). While we focused our evaluation on P/LP variants given their direct relevance to disease, we note that AutoGVP resolved a large number of InterVar’s VUS calls into P, LP, B, LB (**Figure 1C**). Using AutoGVP without the specification of the MedGen Concept ID, we identified 353 P variants and 310 LP variants (total: 663 P/LP variants). Applying Concept IDs associated with cancer, we identified the same 353 P variants when either the ‘--latest’ or ‘--most_ severe’ options were used, but 311 LP variants for ‘--most_severe’ and 310 LP variants for ‘--latest’. One *MEN1* variant (https://www.ncbi.nlm.nih.gov/clinvar/variation/305302/) was classified as LP in “most severe” but classified as VUS using latest or omitting Concept IDs. In contrast, 353 P and 366 LP variants (total: 719 P/LP variants) were identified using InterVar only.

Despite the 56-variant difference in total P/LP count between AutoGVP vs. InterVar, when comparing at the variant level across P/LP/VUS/LB/B classifications, a total of 6,770 variants were discordant. 112 variants were P/LP by AutoGVP, but VUS/LB/B by InterVar. All 112 variants were found in ClinVar: 35 variants with more than 2-stars and 77 1-star classifications. Of the 1-star classifications, 37 came from a single submitter and 40 had conflicting submissions, which were resolved using either “consensus” or the “most recent evaluated date” (**Figure 1B**). Conversely, there were 168 variants P/LP by InterVar, but VUS/LB/B by AutoGVP. Of those 168 variants, 91 were classified by ClinVar and 77 were classified by InterVar. Of the InterVar-classified variants, 76 were downgraded by AutoGVP due to PVS1 adjustments, and one variant that still retained PVS1=1 was demoted to VUS due to removal of PP5, highlighting the over-calling of P/LP variants by InterVar alone.

## 4. Conclusion

Currently available germline sequence variant classification tools are limited by 1) the number of variants that can be classified, 2) the ability to classify only previously reported variants, 3) the ACMG-AMP rules being modified after the time of tool development, 4) the lack of integration with ClinVar, the most extensive and curated germline variant database, and 5) paywall restrictions. Therefore, we developed AutoGVP, an open-source dockerized R workflow that automatically integrates ClinVar and modified InterVar variant annotations with updated ACMG-AMP criteria to classify the pathogenicity of germline sequence variants for research applications. It is important to note that the AutoGVP modifications of InterVar reported here do *not* incorporate gene-specific classification rules developed by ClinGen VCEP. Thus, review of variant classification for genes that have undergone the VCEP process is highly recommended. AutoGVP can be applied to disease-agnostic germline datasets in a fully automated fashion. With the increasing availability of germline exome and genome sequencing data, we anticipate AutoGVP will become a valuable resource for the research community.

## Acknowledgements

This work was supported by National Institutes of Health [R03CA230366 to S.J.D., U2CHL138346 to A.C.R.], the Intramural Research program of the Division of Cancer Epidemiology and Genetics, National Cancer Institute and the Division of Neurosurgery at the Children’s Hospital of Philadelphia. This work was also funded by Gabriella Miller Kids First pilot funds and in part, utilized the computational resources of the NIH HPC Biowulf cluster (http://hpc.nih.gov).

## Author Contributions

Conceptualization: JK, ZV, DRS, SJD, JLR

Methodology: JK, ZV, ASN, RSK, RJC, JLR, DRS, SJD, MAB

Software: ASN, RJC, JK, JLR, MAB, DPM, SP, SJD, RSK

Validation: JK, ASN, JLR, ZG, RJC, MAB, RJC, RSK, ZV, SJD, DRS, SP

Formal Analysis: JK, ASN, RJC

Investigation: JK, ASN, MAB, RJC, SJD, ZV, DRS, RSK

Resources: ACR, PBS, DRS, SJD Data Curation: JK, ASN

Writing - Original Draft: JK, ASN, JLR, DRS, SJD, RJC

Writing - Review & Editing: JLR, SJD, DRS, ZV, MAB, DPM, ZG, PBS, ACR, SP, RSK

Visualization: JK, JLR, RJC Supervision: JLR, DRS, SJD

Project Administration: JLR, SJD, DRS

Funding Acquisition: JLR, DRS, SJD, ACR, PBS

